# Predicting the placement of biomolecular structures on AFM substrates based on electrostatic interactions

**DOI:** 10.1101/2023.06.13.544749

**Authors:** Romain Amyot, Noriyuki Kodera, Holger Flechsig

**Affiliations:** Nano Life Science Institute (WPI-NanoLSI), Kanazawa University, Kakuma-machi, Kanazawa, Ishikawa 920-1192, Japan

## Abstract

Atomic force microscopy (AFM) and high-speed AFM allow direct observation of biomolecular structures and their functional dynamics. Based on scanning the molecular surface of a sample deposited on a supporting substrate by a probing tip, topographic images of its dynamic shape are obtained. Critical to successful AFM observations is a balance between immobilization of the sample while avoiding too strong perturbations of its functional conformational dynamics. Since the sample placement on the supporting substrate cannot be directly controlled in experiments, the relative orientation is *a priori* unknown, and, due to limitations in the spatial resolution of images, difficult to infer from *a posteriori* analysis. We present a method to predict the macromolecular placement of samples based on electrostatic interactions with the AFM substrate and demonstrate applications to HS-AFM observations of the Cas9 endonuclease, an aptamer-protein complex, and the ClpB molecular chaperone. The model also allows predictions of imaging stability under tip-scanning. We implemented the developed method within the freely available BioAFMviewer software package. Therefore, predictions based on available structural data can be made even prior to an actual experiment.

## Introduction

High-speed atomic force microscopy (HS-AFM) allows direct observation of biomolecules during their operation under near-physiological conditions [1,2], with its applications having significantly advanced the understanding of biological processes at the nanoscale [3]. Furthermore, by the combination of HS-AFM and computational modeling even atomistic details of protein function can be inferred [4].

An AFM experiment requires the biological sample to be first deposited on a supporting surface, after which scanning of the molecular surface by a probing tip proceeds to record a topographic image of its shape at a spatial resolution of ∼1-2 nm in the lateral direction and typically less than 0.5 nm in the vertical direction. It is important to understand that the observation of single proteins under HS-AFM is a delicate balance between immobilizing the structure on the supporting surface while at the same time preventing too strong perturbations by immobilization. That is, stable and steady scanning of the protein by the probing tip requires sufficient fixation on the surface through molecular interactions. However, the reliable observation of protein activity rests on the assumption that such interactions (which are not present under physiological conditions or in vitro experiments) do not significantly interfere with the functional conformational dynamics of the protein.

The process of placing a biomolecular sample on the supporting surface and controlling its proper attachment is a challenge at the very start of every HS-AFM observation. Mica, silicon and highly oriented pyrolytic graphite (HOPG) are often used as the supporting substrates. Because of its surface flatness at the atomic level over a large area and easy to prepare surface modifications, the negatively charged Mica substrate is most frequently used. It is possible to modify Mica with specific molecules (e.g., 3-Aminopropyltriethoxysilane (APTES), poly-L-lysin (PLL) or lipid-bilayers), hence altering the charge properties. Furthermore, by the chemical composition of the buffer interactions between the sample and substrate can be modified. Such surface modifications are often critical for successful AFM observations of protein structures and their functional motions [5-7].

It would clearly be valuable to have methods available that can predict the placement of biomolecular structures on the supporting substrate even prior to an AFM experiment being performed, and, on the other side, to facilitate the post-experimental analysis of recorded images to better understand measured AFM topographies. We report here the development of a computational framework to address such issues based on an electrostatic interaction model and demonstrate various applications. The method is implemented within the freely available BioAFMviewer software package [8].

## Results

We illustrate our approach by first considering an idealized situation in which the sample is viewed as a perfectly spherical solid object. Two cases are distinguished. In one case point-like unit charges q_i_ with randomly picked sign are distributed uniformly on the surface of the sphere, while in the second case the two hemispheres carry opposite charges creating a *Janus sphere* (Fig. 1A,B). The AFM supporting substrate is modelled as a 2D solid plate which has point-like charges placed along a regular grid. Details are described in the Methods section.

**Fig. 1:**
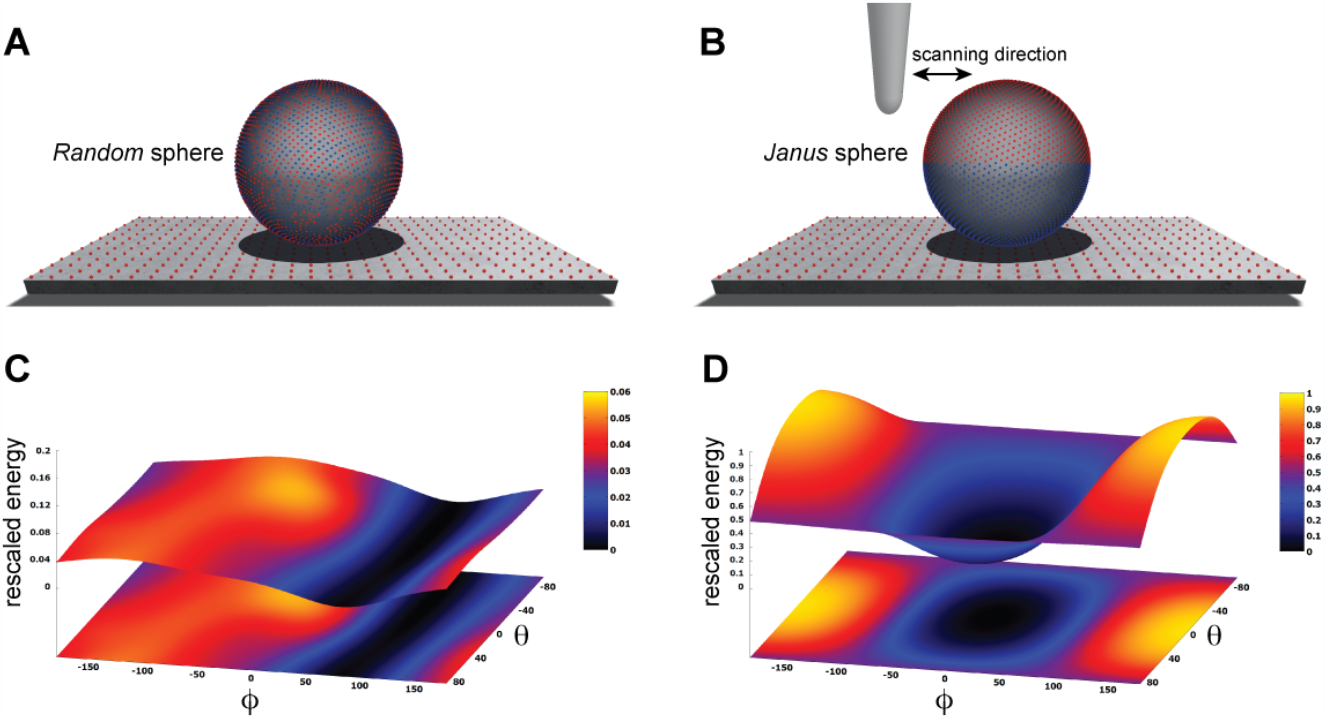
Idealized toy samples. Sphere with 5000 surface point charges of randomly picked sign (A) versus a *Janus* sphere carrying opposite charges separated on either hemisphere (B), each placed on a 2D substrate plate which has point charges arranged along a regular grid. Blue and red colors represent positive and negative unit charges, respectively. The landscape of electrostatic interactions energies for the *random* sphere (C) and *Janus* sphere (D), respectively. In both plots a common energy scale was used by rescaling.

Electrostatic interactions between the sample and the substrate can be described by the Debye-Hückel potential

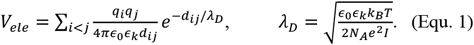

which represents Coulomb interactions of charges *q*_*i*_ and *q*_*j*_ separated by the spatial distance *d*_*ij*_, effectively screened over the Debye length λ_D_. The Debye length can vary between 2.1 nm for ionic strength *I*=20 mM and 0.8 nm *I*=150 mM (ionic strength under physiological conditions). Under such interactions an equilibrium configuration of the sample-substrate complex can in principle be identified as the global minimum in the high-dimensional landscape of all electrostatic energies *V*_*ele*_.

Here, we employ a simplified description resting on the approximation that the sample is placed on top of the AFM substrate (Fig. 1) and its atomistic structure does not undergo any internal conformational changes. We then systematically explore molecular orientations of the sample relative to the substrate by performing rigid-body rotations in 3D space, recording the electrostatic interaction energy for each instantaneous configuration (see Methods). Thus, a landscape of electrostatic interaction energies in the space of appropriately chosen coordinates can be constructed, which shall allow an interpretation of the stability of sample-substrate interactions. We employed the latitude and longitude angles ∅, *θ* to characterize the sample orientation in 3D space (see Methods). A landscape with multiple minima separated by shallow barriers would indicate rather unstable placement of the sample on the surface. This situation is demonstrated for the case of the randomly charged sphere (Fig. 1C) and its interpretation is that a plethora of possible molecular orientations with respect to the stage are practically as likely while a single stable configuration cannot be formed.

The situation is very much different for the Janus sphere, where the landscape shows a highly symmetric shape of a funnel leading into a deep valley with a global minimum that corresponds to a single most stable configuration (Fig. 1D). In this arrangement, the positive charged hemisphere is aligned towards the negatively charged mica substrate contacting it around the pole, and the negatively charged hemisphere is pointed upwards. Deviations from this stable state correspond to uphill motions in the energy landscape, which require forces *F*_*ϕ*_ = *−∂V /∂ϕ* and *F*_*ϕ*_ = *−∂V /∂ϕ*. Since changes in the angle *ϕ* correspond to a rotation of the sample around an axis within the supporting substrate, the component *F*_*ϕ*_ has an intuitive meaning of the force that would be caused by the AFM tip in the horizontal scanning direction (see Fig. 1B).

Proceeding with applications to biomolecular structures, we have first applied a method to compute the electrostatic potential on the molecular surface based on the Coulomb contributions of all amino acids (see Methods). For a given structure, we thus obtained a graphical representation of its molecular surface where values of the electrostatic potential are mapped on a color scale (Fig. 2). Similar graphical representations are typically provided by standard molecular viewers such as ChimeraX [9], Pymol [10] and others.

**Fig. 2:**
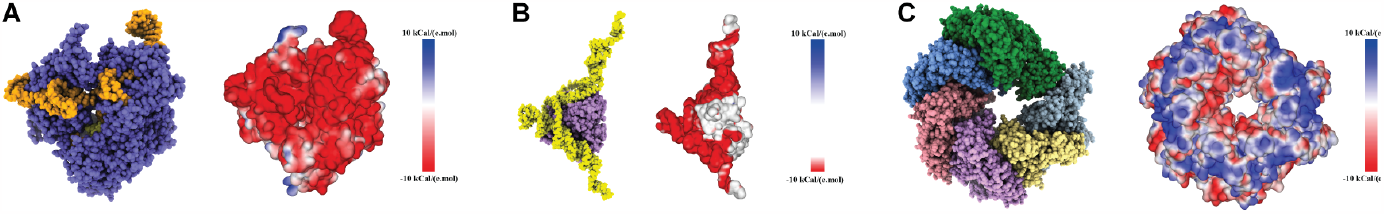
Protein surface electrostatic potential. (A) Molecular structure of the Cas9-RNA-DNA endonuclease complex (left, PDB 4OO8) and the computed surface representation with a coloration representing electrostatic potential values (right). (B) Molecular structure of the aptamer-Cyp24A protein complex (left) and the colored surface representation (right). (C) Molecular structure of the Hsp104 hexamer (left, PDB 5KNE) and the colored surface representation (right).

We have considered three different examples of proteins, a Cas9 endonuclease, an aptamer-protein complex, and the molecular chaperone ClpB. For all cases we have previously applied simulation atomic force microscopy and automatized rigid-body fitting within the BioAFMviewer software package to predict the molecular orientation from resolution-limited HS-AFM topographies [8,11,12], which allowed to disambiguate the arrangement of functional domains and to identify the relative orientation of domains with respect to bound nucleic acids.

Here, we now apply the electrostatic interaction model to predict the 3D molecular placement of the sample on an AFM substrate prior to an actual experiment being performed. We also discuss predictions for the stability of observations. To validate model predictions, we furthermore provide comparison to images from HS-AFM experiments.

### Cas9-RNA-DNA complex

We first considered the Cas 9 endonuclease protein which binds guide RNA and cleaves duplex target DNA with a sequence complementary to the RNA guide, playing a key role in genetic engineering applications (CRISPR-Cas9 genome editing). Several PDB structures of Cas9 complexes are available. Fig. 2A shows the atomistic structure of Cas9-RNA with a bound single-strand target DNA together with the computed molecular surface representation colored according to the computed electrostatic potential. The presence of nucleic acid strands with the phosphate groups in nucleotides generates a negatively charged molecular surface. HS-AFM experiments to visualize structural dynamics of the Cas9-RNA-DNA complex [13] were therefore performed on a modified surface known as APTES-mica which has a positive charge distribution (see Methods). Fig. 3A shows the top two molecular placements on the surface predicted from our electrostatic interaction model. As can be seen from the bottom view perspective, the Cas9 complex binds to APTES-mica with the flat molecular surface that has the negatively charges guide RNA strand attached, which acts like a glue between Cas9 and APTES-mica (Fig. 3C). It should also be noted that the two predicted orientations are related by rotation around the axis perpendicular to the APTES-mica surface. This can be explained as a general phenomenon in the model. For a biomolecule with a typical size of several to tens of nanometers placed on a surface with charges homogeneously distributed along a grid of sub-nanometer spacing, it should be expected that for a given molecular placement any orientations related to rotation around the surface perpendicular axis will result in similar electrostatic interactions patterns and hence be practically indistinguishable.

**Fig. 3:**
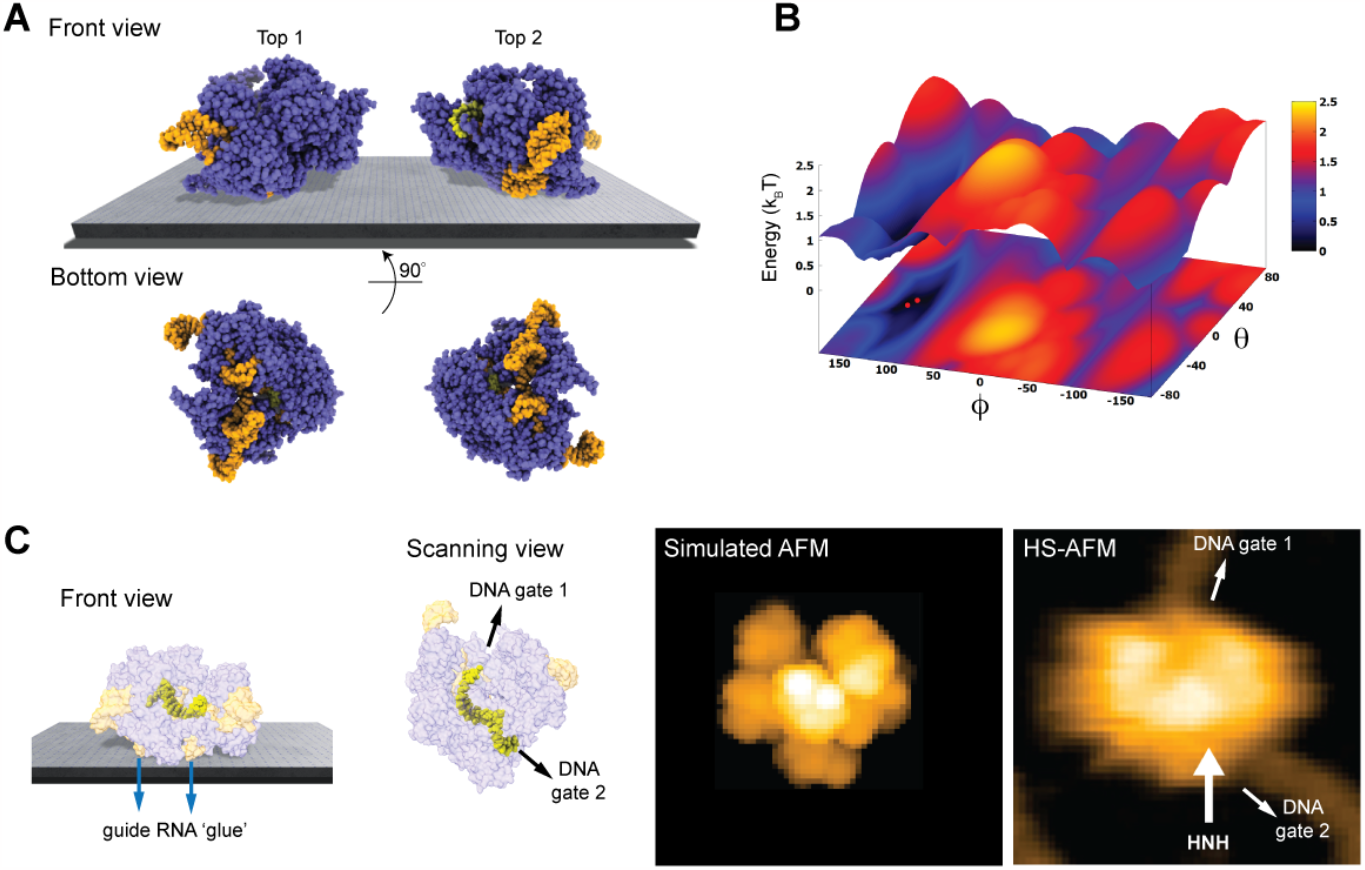
Cas9-RNA-DNA complex. (A) The top two predicted orientations of the protein complex shown on the supporting surface in front view. Additionally, the bottom view perspective displays the structure facing the surface. (B) The landscape of electrostatic interactions energies. Red dots mark locations of the predicted orientations. (C) Left: predicted placement of the Cas9 complex highlighting the gluing role of guide RNA and the parallel orientation of the bound target DNA strand within the Cas9 protein relative to the supporting surface. For better visualization of the DNA located inside the protein, the Cas9 structure is shown in transparent. Middle: The scanning view perspective indicating the position of the two target-DNA gates together with the corresponding simulated AFM topography. Right: HS-AFM image of Cas9 complex with the DNA strand observed at locations very similar to predicted gates (adapted from Ref. [13]).

Looking closer at the predicted orientation of the Cas9 complex relative to the APTES-mica surface, an interesting observation can be made. The bound target DNA strand is located in a tunnel within the Cas9 structure and both the entry and exit paths are not blocked by contacts with APTES-mica.

In fact, the orientation of both DNA gates is roughly parallel to the surface (Fig. 3C). We then compared a simulated AFM image of the molecular structure obtained with the BioAFMviewer software with a snapshot obtained from HS-AFM imaging the dynamics of Cas9 interactions with DNA [13]. As we find, in the experimental image the orientation of the DNA strand in the Cas9 complex correlates remarkably well with the position of the two DNA gates in the predicted molecular orientation. It should also be noted that the predicted molecular orientation of Cas9 relative to the AFM surface based on electrostatic modelling agrees remarkably well with our previous result [8], where automatized rigid-body fitting of the Cas9 structure without the nucleic acids was employed to validate the domain arrangement seen in HS-AFM imaging (see SI material).

Fig. 3B shows the landscape of electrostatic interaction energies in the space of latitude and longitude angles ∅, *θ* which characterize the protein orientation in 3D space. The landscape shows a clear valley localized around the minima which corresponds to the predicted favorable placements of the Cas9-RNA-DNA structural template. This valley is confined by steep walls characterized by the gradient *∂V/∂ϕ* which corresponds to the force magnitude of a perturbation that would be required to destabilize the placement on the protein on the APTES-mica surface. Hence, the presence of such a barrier would resist possible perturbations applied by the AFM tip in the horizontal scanning direction. However, the model simplifications underlying our predictions (see Discussion) do not allow to provide quantitative estimates that could be compared with those obtained from HS-AFM experiments.

Nonetheless, our findings based on electrostatics offer an explanation why under HS-AFM observations functional relative motions of target DNA and Cas9 can be reliably observed, and DNA cleavage could be captured at the single molecule level [13].

### DNA-aptamer protein complex

Next, we considered a complex of a 70-nucleotide DNA aptamer and the CYP24 protein, which has been demonstrated to be relevant for antiproliferative activity in cancer cells and was previously observed under HS-AFM [14]. The 3D atomistic structure of the complex as predicted from molecular docking simulations and the computed molecular surface representation with charge coloring according to the electrostatic potential are shown in Fig. 2B. Similar to the previous case of the Cas9-RNA-DNA complex, the presence of the DNA aptamer which due to the phosphate groups in nucleotides is negatively charged, does generally not allow stable AFM observations on the standard negatively charged bare mica surface. Therefore, the experiment was conducted with a mica surface modified by a positively charged lipid bilayer which self-assembled on its top (for details see [14]).

Taking into account the charge distribution of this specific surface (see Methods) and electrostatic interactions with the aptamer-protein complex, we predict favorable molecular placements a priori to any experiment being performed. Fig. 3A shows the top two candidates found from scanning the space of possible rigid-body orientations relative to the surface. As can be seen in the front view perspective, and particularly well from the bottom view, the predicted orientations are those in which the two longer DNA strands are placed on the surface in a flat arrangement with the CYP24 protein sitting on top. The two predicted orientations are related by rotation around the axis perpendicular to the modified mica surface, and thus being roughly equivalent.

Fig. 3B shows the landscape of electrostatic interaction energies between the aptamer-protein complex and the modified mica surface in the space of latitude and longitude angles ∅, *θ*. It shows a highly localized valley which is confined by steep and high walls and at its bottom has energy minima that correspond to the well-defined placement of the protein-DNA complex predicted from them. The existence of such a highly confined deep energy valley is clearly due to the presence of the DNA aptamer, dominating the electrostatic interactions by attraction forces with the positively charged lipid bilayer on mica. Any structural orientations deviating from the predicted highly stable conformation would be practically impossible, which is confirmed by single molecule HS-AFM observations of the CYP24-aptamer (see SI movie in Ref. [14]).

### ClpB molecular chaperone

As a third application, we chose the ClpB chaperone which is an ATP-powered molecular machine involved e.g. in disaggregation of proteins under heat stress conditions. Functional conformational dynamics of ClpB was previously investigated in HS-AFM experiments [15]. The atomistic structure of the hexameric Hsp104 disaggregase (yeast homologue of bacterial ClpB) in the conformation with bound ATP analog is shown in Fig. 2C. In the chosen orientation the molecular surface representation reveals a ring-shaped region with predominantly positive electrostatic potential. It can therefore be expected that this protein side forms contacts with the mica surface. Our electrostatic interaction model employing a bare mica surface as used in ClpB HS-AFM observations indeed predicted orientations with similar contact surfaces. A single chosen predicted placement is shown in the front view (Fig. 5A). The corresponding bottom view perspective displaying the protein side facing the mica surface together with its molecular surface representation colored according to the electrostatic potential is shown in Fig. 5B. Also shown is the molecular structure in the opposite view corresponding to the scanning perspective together with its surface representation (Fig. 5B). As can clearly be seen, the ring-shaped region with predominantly positive electrostatic potential is guiding the placement of the hexameric chaperone on the negatively charged mica surface.

**Fig. 4:**
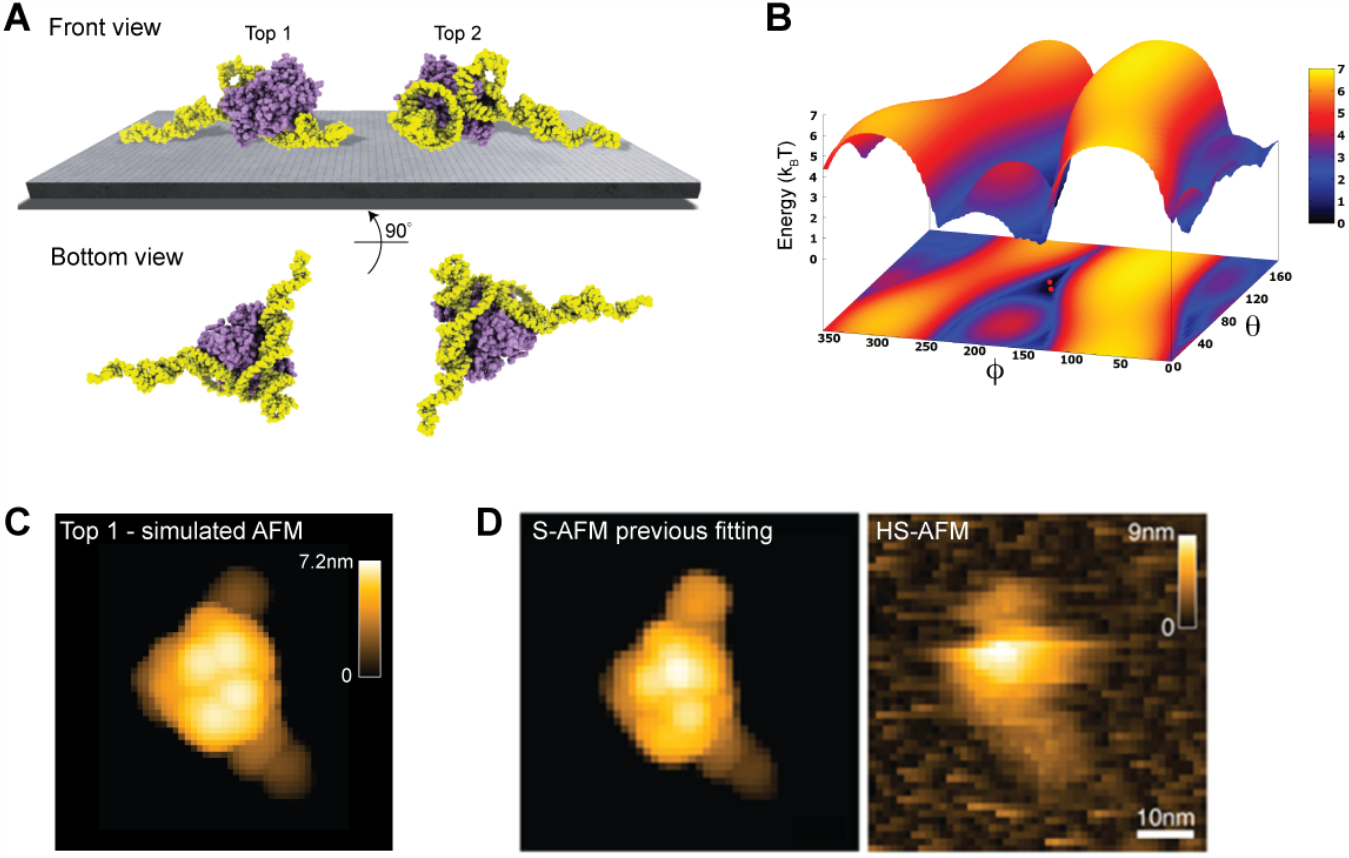
DNA-aptamer CYP24 protein complex. (A) The top two predicted orientations of the protein complex shown on the supporting surface in front view. Additionally, the bottom view perspective displays the structure facing the surface. (B) The landscape of electrostatic interactions energies. Red dots mark locations of the predicted orientations. (C) Simulated AFM image of the top predicted orientation. (D) Simulated AFM image of the molecular orientation (left), identified from previous fitting to a HS-AFM target image (right) based on exhaustive search (images adopted from Ref. [14] with permission, Copyright 2022 American Chemical Society).

**Fig. 5:**
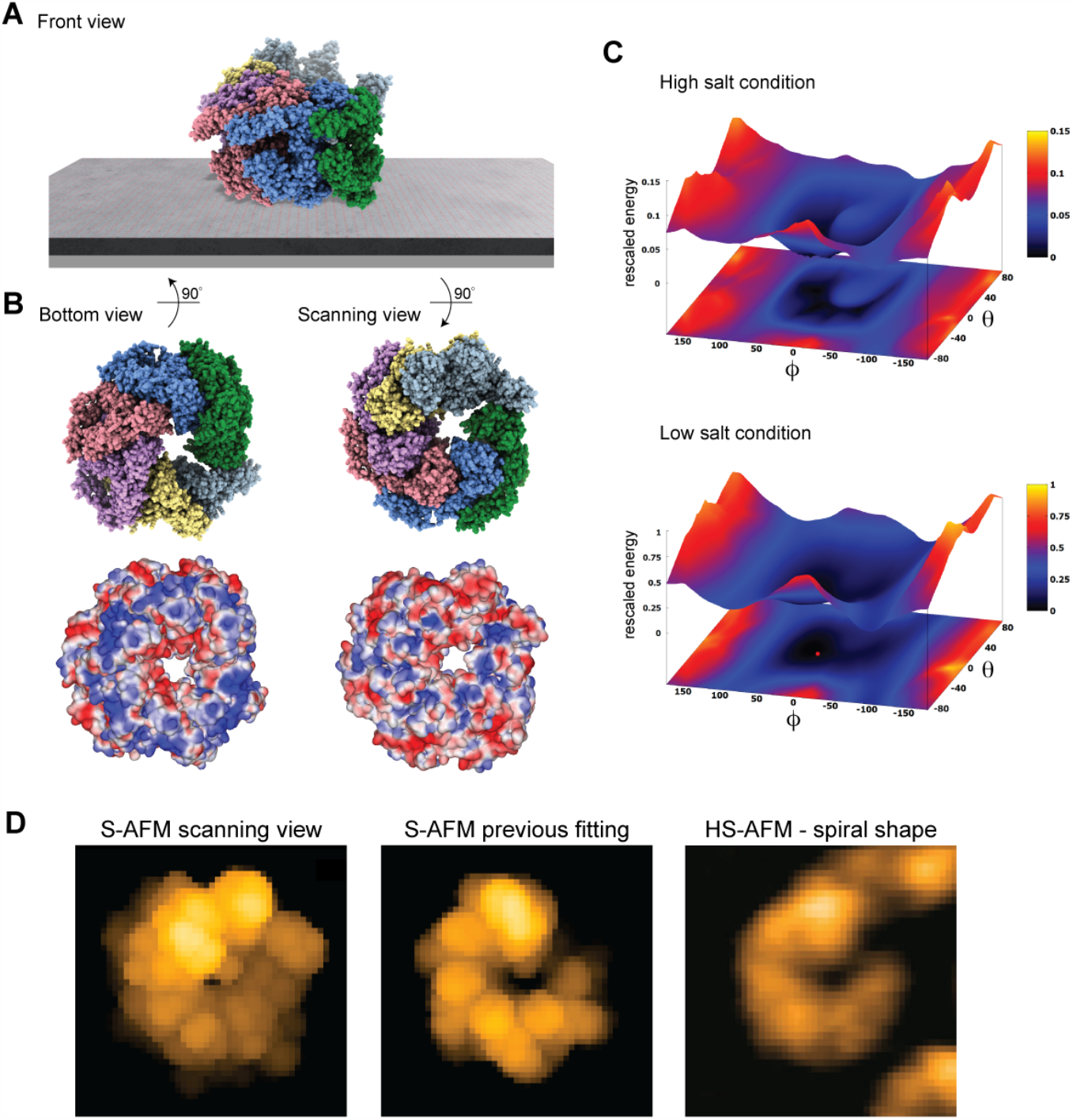
ClpB molecular chaperone. (A) Predicted orientation of the Hsp104 protein structure shown on the supporting surface in front view. (B) Bottom view perspective displaying the atomic structure facing the surface and the corresponding molecular surface representation with colors indicating electrostatic potential values (left). Additionally, the scanning view perspective is provided (right). (C) The landscape of electrostatic interactions energies computed for high and low salt buffer conditions (top and bottom, respectively). For the latter case the red dot marks the location of the predicted orientation. In both plots a common energy scale was used by rescaling.(D) Simulated AFM image of the predicted orientation (left). Simulated AFM image of the molecular orientation (middle), identified from previous fitting to a HS-AFM target image (right, taken from Ref. [15]) based on exhaustive search.

An interesting aspect is that for successful imaging of ClpB under HS-AFM the buffer composition was critical as stated in Ref. [15]: *“The salt concentration was a key to successful imaging of TClpB because high salt concentrations such as 150 mM KCl weakened the affinity of molecules to mica substrate, resulting in fast diffusion of molecules and thus hampering imaging. Therefore, we used a lower concentration of KCl (20 mM) which enabled moderate binding of ClpB onto mica substrate*.*”* While in the employed electrostatic model the buffer conditions can only be phenomenologically accounted for by the parameter for the ionic strength, our predictions can still provide an explanation of this situation. In Fig. 5C we show the landscape of electrostatic interaction energies between the Hsp104 structural template and the mica surface for two cases, corresponding to high- and low-salt concentrations, respectively. Common to both landscapes is the presence of a valley isolating favorable protein placements on mica. However, the stark difference between them is that under low-salt conditions, electrostatic interaction energies, and therefore the barriers around the valley, are larger by one order of magnitude as compared to the high-salt case. This is because in the latter case, electrostatic interactions are screened over a much shorter Debye length (see Equ. 1). Hence, the interpretation is that a buffer condition with low-salt concentration significantly stabilizes the formation of electrostatically favorable orientations of ClpB on mica and therefore allows reliable imaging under HS-AFM scanning.

A simulated AFM image of the predicted shown in Fig. 5A,B resembles the spiral shape topography seen in HS-AFM imaging (Fig. 5D), which arises from the domain protrusions in the hexameric arrangement. Interestingly, our previous result of automatized fitting the PDB structural template into the same HS-AFM image predicted the hexamer structure to be in the opposite upside-down orientation. There, fitting was based on exhaustive sampling of possible molecular orientations without an underlying physical model, aiming to identify the orientation whose simulated AFM image best matched to the target HS-AFM image. In fact, the thus obtained simulated AFM image matches much better to the HS-AFM image compared to the one obtained from our electrostatic model predictions (Fig. 5D). A well-known drawback in the interpretation of results is that simulated topographies (like the measured AFM topographies) have a limited spatial resolution. Especially for symmetrically shaped proteins this may lead to ambiguities. While the atomistic structure on opposite sides of the ClpB ring is clearly distinct, the corresponding simulated AFM images resulting from a convolution of the tip shape with the molecular structure can show similar looking spiral shapes.

Predictions based on our electrostatic model should in principle be prioritized over the sampling method without any physical interactions. A so far overlooked issue is that HS-AFM observations were performed under a high protein concentration imaging assembly of ClpB rather than single molecules. Therefore, additional inter-molecular interactions may influence the placement on the mica substrate and imaging stability.

## Discussion

We addressed a simple question relevant in all biomolecular scanning probe experiments: can the sample placement on the supporting substrate be predicted? – with the answer obviously being, of course. Our approach based on electrostatic interactions allows such predictions considering available structural data prior to an actual experiment. We demonstrated its validity in applications to HS-AFM imaging to not only confirm resolution-limited imaging results using atomistic-level information, but also to offer an explanation about the stability of observations. Buffer conditions are considered in the model by phenomenologically including e.g. the salt concentration via the ionic strength in the Debye-Hückel form of interactions, and the pH value affecting the biomolecular surface electrostatic potential.

Our model predictions rest on the gross simplification of neglecting any conformational motions within the biomolecule which certainly play a role in the interactions with the AFM substrate. In that sense, the presented approach builds on our previous work on rigid-body sampling to infer atomistic structure from AFM images by automatized fitting [11]. As we demonstrated, efficient predictions which agree remarkably well with experimental observations can be obtained despite such approximations. While molecular modeling to resolve conformational changes is certainly possible, its implementation would drastically increase the computational expense for predictions.

The developed methods are implemented in our BioAFMviewer package freely available at www.bioafmviewer.com, allowing for convenient applications within a well-established user-friendly interactive software interface. We are inviting the BioAFM community to use the new tool and are anticipating constructive feedback.

## Methods

### Substrate modelling

The AFM substrate is modelled as a plane which has point-like charges *q* placed along a regular grid with spacing parameter *d*. For the mica surface (charge density −*e/*0.48nm^2^ [M1]) we use *q* = −*e* and 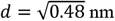. For the lipid bilayer surface (charge density +*e/*0.26nm^2^ for a DPPC lipid composition [M2]) we use *q* = *+e* and 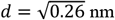. The *Electrostatics Application* within the BioAFMviewer allows to modify such parameters to consider other AFM substrates too.

### Electrostatic potential calculation

To construct the 3D surface of a macromolecular structure, the well-known *marching cubes* discretization algorithm [M3] was employed, representing it by a set of triangles used for graphical rendering. For each triangle the electrostatic potential is evaluated at the center of mass (vertex) as the sum of Coulomb potentials arising from partial charges of all atoms. Hence, the potential for vertex *i* is *V*_*i*_ = (4*π* _0_)^−1^Σ_*j*_ *q*_*j*_*/d*_*ij*_, where *q*_*j*_ is the partial charge of atom *j, d*_*ij*_ is the distance between vertex *i* and atom *j*, and *ϵ*_0_ is the vacuum permittivity. This allows to compute the surface electrostatic potential for a given PDB structure. The *Electrostatics Application* within the BioAFMviewer implements a graphical representation of the generated 3D molecular surface where values of the electrostatic potential are visualized via a color scale. In the applications for Cas9, aptamer-CYP24 and ClpB, the partial charges of atoms were computed at pH value 7.0 condition.

### Orientation sampling, energy landscape and prediction

For sampling 3D rigid-body orientations of a biomolecular structure, we discretized the search space evenly using the Fibonacci lattice algorithm [M4, M5]. In the applications for Cas9, aptamer-CYP24 and ClpB, sampling was performed for a set of 2000 conformations. The *Electrostatics Application* within the BioAFMviewer gives a choice for this number. For each single orientation, direct contact to the AFM substrate was always assumed, and the electrostatic interaction energy between all substrate point charges and the biomolecular surface was computed according to the Debye-Hückel form. For a single substrate point charge *q*_*i*_ the electrostatic potential energy is 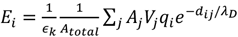, where *V*_*j*_ is the electrostatic potential value of triangle *j* (with area *A*_*j*_) on the sample surface, *A*_*total*_ is the total sample surface area, *d*_*ij*_ is the distance between the triangle center of mass and the charge *q*_*i*_, and *ϵ*_*k*_ is the dimensionless relative permittivity. Area weights were introduced because the *marching cubes* algorithm does not discretize the sample surface evenly, resulting in a heterogeneous density of vertices (triangle center of mass). To remove the density dependence in the electrostatic potential energy, individual contributions of triangles were therefore weighted considering their fraction to the total surface area. After completed sampling, a visualization of the energy landscape in the space of latitude and longitude angles ∅, *θ* (characterizing the sample orientation relative to the substrate) was obtained. For presentation purposes, we shift the energy scale such that the global minimum has zero energy. The prediction of most favorable placements on the substrate was based on identifying the minima of the landscape. In the software the top five candidates, corresponding to the five lowest values, are displayed.

### BioAFMviewer workflow

The developed methods are implemented via the *Electrostatics Application* tool within the BioAFMviewer interactive software interface. To use this application, the user has to upload a PQR file of the biomolecular structure which, as compared to a regular PDB file, contains information about partial charges of atoms. For a given biomolecular structure such data can be conveniently obtained using for example the PDB2PQR application [M6] (server https://server.poissonboltzmann.org/pdb2pqr), which also considers calculations at variable pH value and for different force fields. After loading the PQR file, the biomolecular structure can be displayed in the surface representation with coloring according to the calculated electrostatic surface potential. In the *Electrostatics Application* tool, the user can either choose the AFM substrate from a list of commonly employed examples with preset charge distribution or provide alternative values. The prediction of electrostatically favorable biomolecular orientations can be started after fixing the size of the sampling set. After completed sampling, the landscape of electrostatic interaction energies is displayed and the five most favorable molecular placements on the substrate can be visualized. Other orientations can be retrieved from the interactive landscape.

### Simulation atomic force microscopy

We have employed simulation AFM to compare the results from electrostatic predictions with measured HS-AFM topographies. Simulation AFM computationally emulates AFM scanning to convert available biomolecular structures into simulated AFM images that can be correlated with experimentally obtained images. It is based on the non-elastic collisions of a rigid cone-shaped tip with a rigid Van-der-Waals sphere atomistic model of the biomolecular structure. For details we refer to our previous work [7]. Simulation AFM calculations were performed within the BioAFMviewer software platform [7,10].

